# Ligand-independent EGFR oligomers do not rely on the active state asymmetric kinase dimer

**DOI:** 10.1101/2020.04.24.056077

**Authors:** Patrick O. Byrne, Kalina Hristova, Daniel J. Leahy

## Abstract

The human Epidermal Growth Factor Receptor (EGFR/ERBB1) is a Receptor Tyrosine Kinase (RTK) that forms active oligomers in response to ligand. Much evidence indicates that EGFR/ERBB1 forms oligomers in the absence of ligand, but the structure and physiological role of these ligand-independent dimers remain unclear. We use fluorescence microscopy to measure the oligomer stability and FRET efficiency for homo- and hetero-oligomers of fluorescent-protein labeled forms of EGFR and its paralog, Human Epidermal Growth Factor Receptor 2 (HER2/ERBB2) in vesicles derived from native cell membranes. Both receptors form ligand-independent oligomers at physiological plasma membrane concentrations. Mutations introduced in the EGFR kinase region at a key interface within the active state dimer alter the FRET efficiency within ligand-independent EGFR oligomers but do not affect their stability. These results indicate that ligand-independent EGFR oligomers do not require this interface and that the inactive state ensemble is distinct from the EGFR active state ensemble.

## Introduction

Human Epidermal Growth Factor Receptor (EGFR/ERBB1) and its paralogs Human Epidermal Growth Factor Receptor 2 (HER2/ERBB2), HER3/ERBB3, and HER4/ ERBB4, collectively known as ERBBs, are Receptor Tyrosine Kinases (RTKs) that are essential for normal growth and development (*1*).

Abnormal activation of both EGFR and HER2 is associated with multiple human cancer types, and each is the target of several anticancer agents (*2*). ERBBs are Type I integral membrane proteins composed of an extracellular domain (ECD) that is made up of four distinct subdomains, an alpha-helical transmembrane domain (TMD), and an intracellular domain (ICD) comprising a juxtamembrane region, a tyrosine kinase domain and a nonglobular ~230 amino-acid C-terminal tail. The canonical mechanism by which ERBBs are thought to act is ligand-dependent oligomerization, which results in stimulation of the intracellular kinase, trans-autophosphorylation, recruitment of effector proteins, and initiation of intracellular signaling cascades (*1, 3–5*).

The reported levels of endogenous EGFR expression vary several orders of magnitude depending on cellular context (*6–15*). At least seven studies have reported that a fraction of cell-surface EGFR is dimeric in the absence of ligand (hereafter called ‘ligand-independent dimers’) (*16–22*) (Supplementary Table S1), indicating that the receptor concentration in the plasma membrane might specify the ligand-independent oligomeric fraction in accordance with the law of mass action (*23, 24*). Despite much study, the structure and signaling role of ligand-independent dimers remain unclear.

Mutations or deletions in EGFR and HER2 result in ligand-independent receptor activation, indicating that in addition to being activated in the presence of ligand, ERBBs are actively inhibited in the absence of ligand (*25–30*). The mechanism for this autoinhibition remains incompletely understood, however. In contrast, much is known about how ligand binding promotes ErbB activation (*31, 32*). In the absence of ligand, the EGFR, ERBB3, and ERBB4 ECDs adopt a tethered conformation that buries an extended beta-hairpin loop (*33–36*). Binding of ligand to these ECDs stabilizes an extended conformation in which an extended β-hairpin (termed the ‘dimerization arm’) is exposed and mediates formation of an active receptor dimer. In this dimer the intracellular kinase domains adopt an asymmetric dimer interaction in which the C-lobe of a ‘donor’ kinase contacts the N-lobe of a ‘receiver’ kinase, stabilizing the active conformation of the receiver kinase (*37*). Comparatively little is known about the structure and physiological role of either the TMD and the ICD in the inactive state, although mechanisms by which the TMD may function in the inactive state have been proposed (*35, 38, 39*).

Here we report quantitative Förster resonance energy transfer (FRET) measurements of homo- and hetero-interactions between near full-length EGFR and HER2 in vesicles derived from plasma membranes. We confirm previous findings that EGFR and HER2 are in dynamic equilibrium between monomeric, homo-oligomeric and hetero-oligomeric forms at physiological receptor concentrations (*40*). Since we are unable to distinguish dimers from higher-order oligomers (n ≥ 2) by our method we refer to “oligomers” rather than “dimers”, although single-molecule photobleaching and fluorescence intensity fluctuation studies indicate that ligand-independent EGFR oligomers are predominantly dimeric (*21, 23, 41*). We observe that mutations in the asymmetric kinase interface do not affect the stability of ligand-independent EGFR oligomers, indicating that an alternate conformation to the active dimer exists within the ligand-independent oligomer ensemble.

## Materials and Methods

### Plasmid Construction

The coding sequences for variants of EGFR, HER2, EYFP and mCherry were amplified using the polymerase chain reaction and cloned into pCDNA3.1(+) (*Life Technologies*). The dimerization-impaired EYFP-A206K variant was used. The missense variants EGFR-I607Q, EGFR-V948R, EGFR-L858R were generated using site directed mutagenesis.

### Cell culture and transfection

Chinese hamster ovary (CHO) cells and A431 cells were maintained in Dulbecco’s modified Eagle Medium (DMEM F12) supplemented with 2 mM glutamine, 5% FBS and were grown at 37 °C, 5% CO2. For imaging experiments, CHO cells were seeded in 35 mm dishes at a density of 2 x 104 cells per well and grown for 24 hours prior to transfection. Cells were transfected with plasmids using Lipofectamine 3000 (*Life Technologies*) according to the manufacturer’s protocol.

### Generation of monoclonal CHO cell lines expressing EGFR-EYFP

CHO cells stably expressing EGFR-EYFP were generated using the pCDNA3.1(+) manual as a guide. Briefly, cells transfected with EGFR-EYFP in the pCDNA3.1(+) vector were grown for several days in medium without antibiotics. G418 was added to a final concentration of 1 mg/mL and the cells were cultured until individual colonies appeared. All colonies were pooled into a single polyclonal culture, from which monoclonal populations were derived using fluorescence activated cell sorting on a FACSCalibur instrument (BD Biosciences). Resulting cell lines were maintained in DMEM-F12 supplemented with 10% FBS, 1 mg/mL G418, and 25 mM HEPES pH 7.2. Stable expression of EGFR-EYFP in these cell lines was assayed by confocal microscopy and western blot (see below).

### Vesiculation, Image Acquisition and Analysis

All vesiculation procedures were performed as described (*42, 43*). For ligand addition experiments, EGF was added to the vesicles to a final concentration of 100 nM and incubated for at least 1 hour prior to imaging. Vesicle images were acquired using a Nikon C1 laser scanning confocal microscope at 60X magnification (water immersion objective). For each vesicle, three scans were recorded: a ‘donor’ scan (λexc = 488 nm, λem = 500-530 nm), an ‘acceptor’ scan (λexc = 543 nm, λem = 650 nm longpass), and a ‘FRET’ scan (λexc = 488 nm, λem = 565-615 nm). Argon (488 nm) and He-Ne lasers were used as excitation sources. The image pixel dimensions were 512 x 512. The pixel dwell time was 1.68 μs, and the gains were set to 8. The Förster radius for the EYFP-mCherry pair was calculated to be 53.1 Å. All images were processed using a MATLAB program written in the Hristova laboratory which calculates the FRET efficiency for each vesicle. A detailed description of the analysis is found elsewhere (*44, 45*). Absolute protein concentration in the membrane was calculated by comparing the fluorescence intensity in vesicles with the intensities measured from a dilution series of fluorescent protein standards (EYFP and mCherry). Bleed-through coefficients were calculated for each experiment (typically ~0.3 and ~0.2 for EYFP and mCherry, respectively). Where appropriate, the processed data (FRET vs. concentration) were fit to a monomer-dimer equilibrium model using Graphpad Prism:

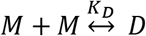

where the dimer association constant, K_D_, is

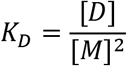

and the total receptor concentration, [*T*] is a function of the concentration of monomers, [*M*], and dimers, *D*]

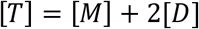

The dimeric fraction for each vesicle, *f_D_*, is given by the equation:

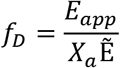

where apparent *E_app_* is the apparent FRET efficiency, *X_a_* is the fraction of acceptor molecules in a given vesicle, and 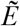 is the FRET value within a dimer.

### EGF Stimulation of ErbB proteins in CHO cells

EGF was expressed in *E. coli* and purified as described (*46*). CHO cells were grown to 90% confluency, then transfected and grown for 14 hours, then serum starved for 7 hours at 37 °C in Ham’s F12 supplemented with 1 mg/mL BSA. The cells were incubated in starvation media in the presence or absence of 100 nM EGF (5 minutes, 37 °C). The medium was aspirated, the cells were washed twice with cold PBS (1X, plus 1 mM Na_3_VO_4_), then lysed for 15-20 minutes at room temperature in RIPA buffer (150 mM NaCl, 50 mM Tris pH 8, 1% NP40, 0.5% w/v sodium deoxycholate, 0.1% sodium dodecyl sulfate, 1 mM Na_3_VO_4_). The lysates were clarified by centrifugation and the total protein concentration of the supernatants was determined by BCA assay (*Pierce, Life Technologies*). Lysate protein concentrations were normalized, and the samples were analyzed by SDS-PAGE and western blot to detect phosphotyrosine (4G10, *Millipore*), EGFR (D38B1), EGFR pY1068, HER2 (29D8), or HER2-pY1221/1222 (all ERBB antibodies purchased from *Cell Signaling Technology*).

## Results

### Quantitative Imaging FRET Microscopy

To enable measurement of homo- and hetero-ErbB interactions in native cell membranes using FRET microscopy, human EGFR and HER2 variants labeled at their C-termini with a fluorescent donor (EYFP) or acceptor (mCherry) protein were expressed in Chinese Hamster Ovary (CHO) cells. ERBBs have ~200 amino-acid disordered regions at their C-termini, which, along with 8 amino-acid Gly-Ser linkers inserted between each ErbB and its fluorescent protein (fp) fusion partner, are presumed to mitigate effects of fluorescent dipole orientation on observed FRET (Supplementary Fig. S1) (*47*). Expression levels of full-length forms of fp-tagged EGFR and HER2 proved too low to characterize interactions with confidence. EGFR and HER2 variants with C-terminal deletions were thus tested to identify minimal deletions that preserve activity and result in improved expression levels. Removal of 107 and 25 residues from the C-termini of EGFR and HER2, respectively, generated the variants EGFR-Δ107-fp and HER2-Δ25-fp and resulted in proteins that express well and are functional as judged by anti-phosphotyrosine Western blots (Fig. 1A).

**Figure 1.**
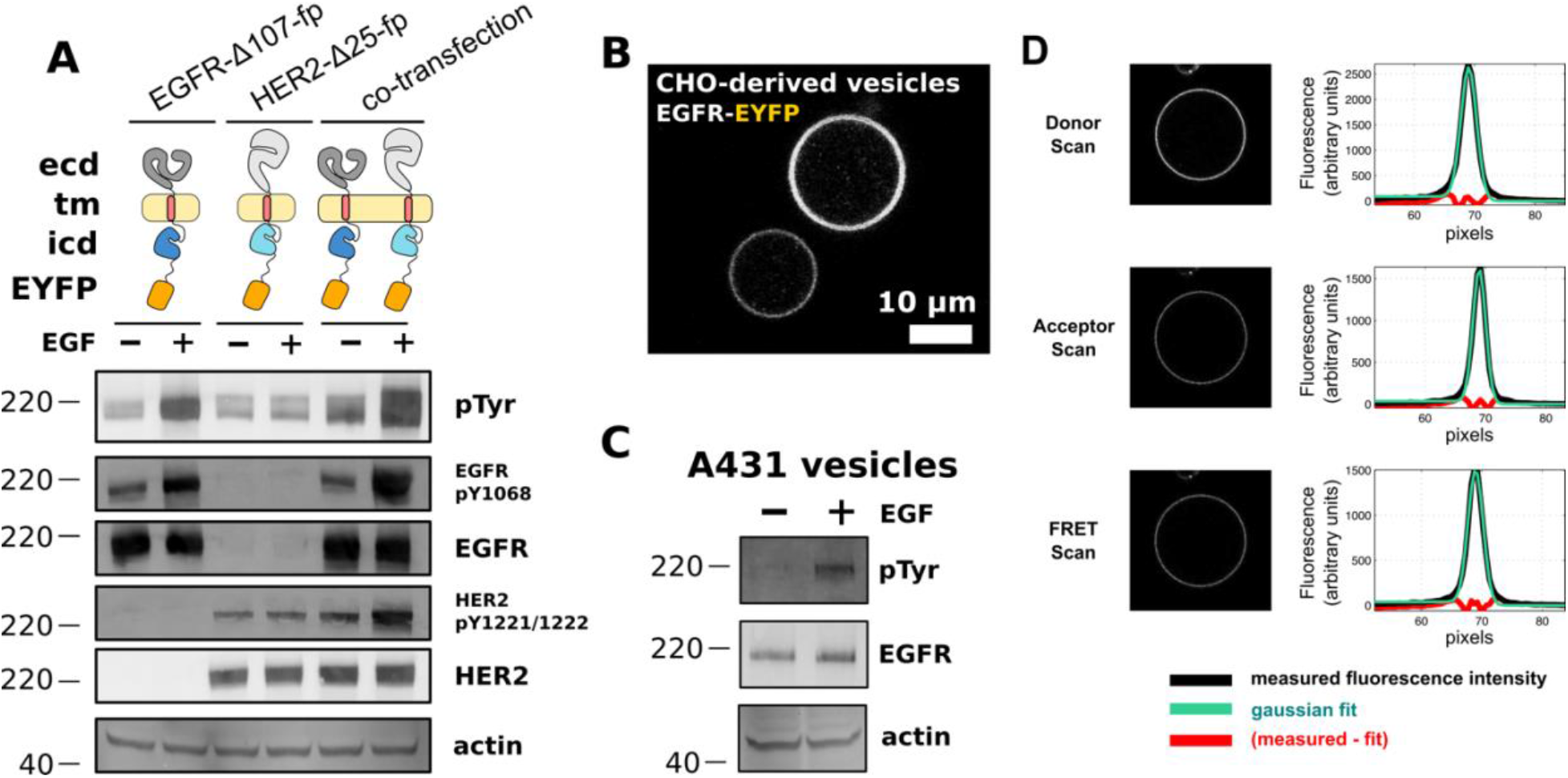
FRET microscopy of near full length EGFR and HER2 in CHO cell vesicles. (A) Western blot analysis of transient expression and EGF-dependent phosphorylation of EGFR-Δ107-FP and HER2-Δ25-FP in CHO cells. Blots are representative of three independent experiments. Primary antibodies are indicated at the right of each blot. ECD = extracellular domain, TM= transmembrane domain, ICD = intracellular domain, pTyr = phosphotyrosine. (B) Confocal image of two vesicles derived from the plasma membranes of CHO cells expressing EGFR-EYFP. Scale bar is 10 μm. (C) Western blot analysis of EGFR in vesicles derived from A431 cells. EGFR becomes phosphorylated in response to EGF in the presence of ATP. (D) Representative scans for a FRET experiment. Plots to the right of each vesicle image show the average fluorescence intensity (y-axis) as a function of radius from the center of the vesicle (x-axis, in units of pixels). Molecular weight markers (kDa) are indicated to the left of each western blot.

Homo- and hetero-interactions between EGFR-Δ107-fp and HER2-Δ25-fp were measured using quantitative imaging FRET (QI-FRET) microscopy (*44, 45, 48*). Briefly, plasma membrane vesicles from CHO cells transiently transfected with fp-labeled EGFR and/or HER2 were generated by incubating live cells in a hyperosmotic solution (Fig. 1B) (*42, 43*). Endogenous EGFR in vesicles derived from A431 cells retains EGF-dependent phosphorylation indicating that vesiculation does not impair ERBB function (Fig. 1C). The concentrations of donor- and acceptor-labeled ERBBs as well as the FRET efficiency between labeled ERBBs in vesicles were obtained from three fluorescence scans (Fig. 1D). Cells in the same transfection experiment display a wide range of receptor expression levels as judged by fluorescence intensity (Fig. 1B), which allows FRET efficiency to be determined as a function of receptor concentration. Following correction of FRET signals for FRET arising from density-dependent proximity (known as ‘proximity’ or ‘bystander’ FRET) (*49*) (Supplementary Figs. S2, S3, S4) and for the relative fraction of donors and acceptors in each vesicle (*44, 48*), apparent two-dimensional dissociation constants can be determined.

The oligomeric state of fluorescently-labeled proteins affects the FRET signal (*50*). The FRET changes we observe fit a monomer-dimer model as well as monomer-dimer-tetramer or monomer-trimer models. We thus report fits to a monomer-dimer equilibrium, although the receptors likely form higher-order oligomers in the presence of EGF (*23, 41*). This fit depends on two parameters: the ‘intrinsic FRET efficiency’, 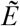, which equals the FRET efficiency within a dimer, and the two-dimensional dissociation constant, *K_d_*, in units of molecules/μm2 (*44, 48*). In cases where the association is strong at all receptor concentrations, we assume that the intrinsic FRET value (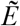) is equal to the mean observed FRET value. As the presence of higher-order oligomers is likely (*41*), especially when ligand is present, we use the term ‘oligomer’ throughout to denote complexes of two or possibly more receptors.

### Near full-length EGFR forms ligand-independent oligomers at physiological plasma-membrane concentrations

The FRET efficiency for EGFR-Δ107-fp increased as a function of concentration and fit well to a monomer-dimer equilibrium model, with a two-dimensional K_d_ of 156 molecules/μm_2_ (Fig. 2A) and a FRET efficiency value of 0.35 +/- 0.06 (Table 1), consistent with previous observations (*23, 24*). Addition of EGF or substitution of the EGFR extracellular domain with a constitutively dimeric immunoglobulin Fc-domain results in constant FRET efficiency values of 0.29 +/-0.03 and 0.29 +/-0.02, respectively, as well as constitutive receptor phosphorylation (Supplementary Fig. S5). The cancer-associated L858R mutation has been shown to promote EGFR oligomerization in the absence of ligand (*51*), and introduction of L858R into EGFR-Δ107-fp results in higher FRET efficiency values over all concentration ranges measured indicating increased homo-dimerization in the absence of ligand (Fig. 2). Observation of higher FRET efficiencies under conditions known to activate EGFR provides confidence that our observations reflect physiological behavior. The FRET efficiency differences become less apparent when the FRET data are binned over all observed receptor concentrations (Figs. 2B, 2D and 2F), underscoring the need for measurements over a range of concentrations.

**Table 1.** Statistics for fits to a monomer-dimer equilibrium model

| Proteins | [EGF]<br>(nM) | $K_{app}$ | (95% CI) | $\tilde{E}$ | (95% CI) |
| --- | --- | --- | --- | --- | --- |
| EGFR | 0 | 156 | (49 to 339) | 0.35 | (0.28 to 0.41) |
| HER2 | 0 | ND* | (ND)# | 0.29 | (0.27 to 0.31) |
| EGFR + HER2 | 0 | ND* | (ND)# | 0.26 | (0.24 to 0.28) |
| EGFR | 100 | ND* | (ND)# | 0.29 | (0.26 to 0.32) |
| EGFR-L858R | 0 | 27 | (0 to 190) | 0.37 | (0.26 to 0.48) |
| IgG-Fc/EGFR chimera | 0 | ND* | (0 to 21.31) | 0.29 | (0.27 to 0.31) |
| EGFR-I706Q | 0 | 65 | (8 to 183) | 0.46 | (0.39 to 0.53) |
| EGFR-V948R | 0 | 364 | (212 to 573) | 0.70 | (0.63 to 0.77) |
| * best-fit value for $K_{app}$ was less than 1 | | | | | |
| # confidence interval was not determined |  |  |  |  |  |

**Figure 2.**
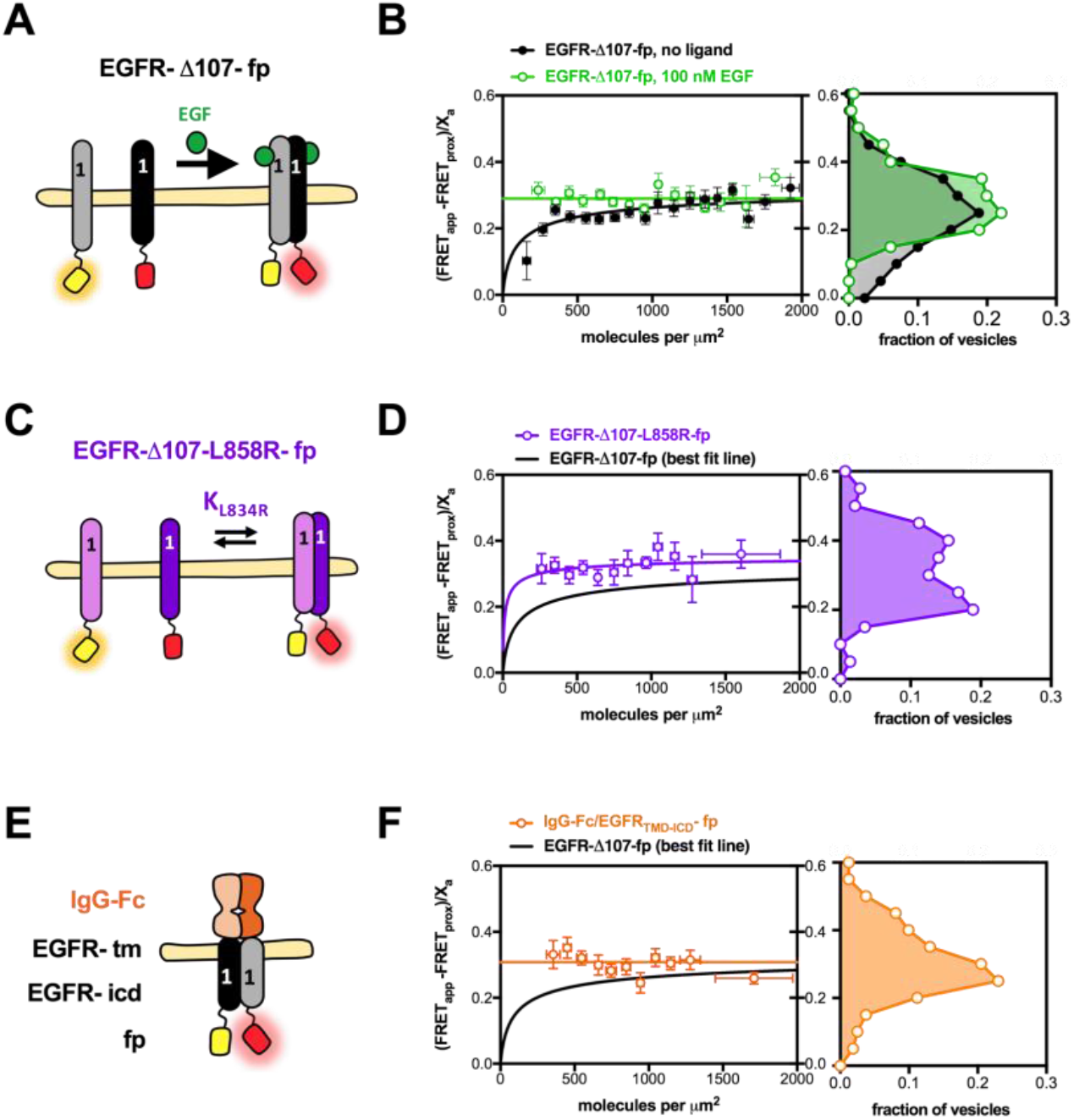
Near full-length EGFR forms ligand-independent oligomers at physiological plasma membrane concentrations. FRET efficiency for EGFR-Δ107-fp homo-oligomers. (A, C and E) Cartoon representations of fluorescent-protein-linked (A) EGFR wildtype, (C) L858R and (E) IgG-Fc/EGFR. (B, D, F) FRET efficiency plots for (B) EGFR-Δ107-fp wildtype, (D) L858R and (F) IgG-Fc/EGFR. Each panel in D, E, and F contains two graphs. The graph on the left shows the FRET efficiency [(FRET_app_ - FRET_prox_)/X_a_)] (y-axis) as a function of concentration (x-axis). The FRET values were projected onto the y-axis and displayed in the graphs to the right to yield the FRET efficiency [(FRET_app_ - FRET_prox_)/X_a_)] as a function of the fraction of all vesicles. FRET_app_ is the apparent FRET efficiency, FRET_prox_ is the theoretical FRET efficiency that results from nonspecific interactions, and X_a_ is the fraction of acceptor molecules in a given vesicle. Binned data points are shown as circles, error bars represent the standard error in x and y. The best fit to a monomer-dimer equilibrium model is represented by solid lines. Panel B shows conditions for EGFR-Δ107-fp in the absence (black) and presence (green) of 100 nM EGF.

### HER2 forms homo- and hetero-oligomers in the absence of ligand

FRET efficiency values for homo-dimerization of HER2-Δ25-fp and associations between HER2-Δ25-fp and EGFR-Δ107-fp were ~0.3 and varied little with concentration, consistent with formation of constitutive ligand-independent HER2 homo-oligomers and EGFR/HER2 hetero-oligomers (Fig. 3, Table 1). These results indicate HER2 self-associates more strongly than EGFR in the absence of ligand.

**Figure 3.**
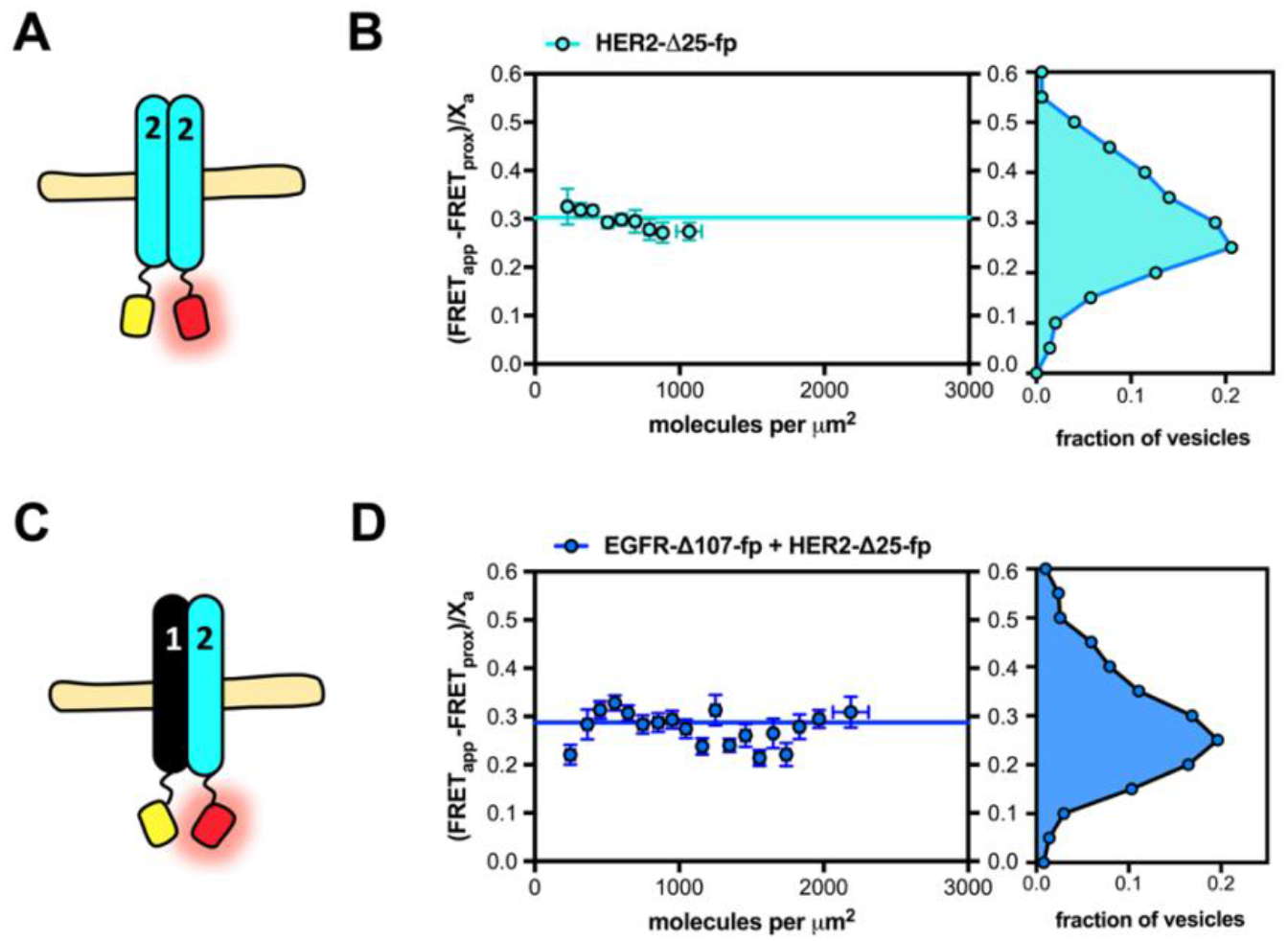
HER2 forms ligand-independent homo-oligomers and hetero-oligomers. FRET efficiency for HER2-Δ25-fp homo-oligomers and hetero-oligomers. (A and C) Cartoon representations of fluorescent-protein-linked (A) HER2-Δ25-fp and (C) HER2-Δ25-fp co-expressed with EGFR-Δ107-fp. (B, D) FRET efficiency plots for (B) HER2-Δ25-fp and (D) HER2-Δ25-fp co-expressed with EGFR-Δ107-fp. Each panel in B and D contains two graphs. The graph on the left shows the FRET efficiency [(FRET_app_ - FRET_prox_)/X_a_)] (y-axis) as a function of concentration (x-axis). The FRET values were projected onto the y-axis and displayed in the graphs to the right to yield the FRET efficiency [(FRET_app_ - FRET_prox_)/X_a_)] as a function of the fraction of all vesicles. FRET_app_ is the apparent FRET efficiency, FRET_prox_ is the theoretical FRET efficiency that results from stochastic, nonspecific interactions, and X_a_ is the fraction of acceptor molecules in a given vesicle.

### EGF-independent EGFR phosphorylation increases with increasing EGFR expression but does not result in pathway activation

Consistent with previous observations, ligand-independent EGFR oligomer formation increases at higher EGFR concentrations, in accordance with the law of mass action (Fig. 2) (*23, 24*). To assess the activity of EGFR oligomers formed in the absence of ligand, we generated a panel of CHO cell lines stably expressing full length EGFR with EYFP at its C-terminus (EGFR-EYFP) and sorted cells to select cell lines expressing increasing EGFR-EYFP levels. The apparent EGFR-EYFP surface concentration of selected cell lines was measured by observing fluorescence in vesicles derived from each cell line. Despite extensive effort, we were unable to isolate a cell line stably expressing EGFR-EYFP at levels higher than ~200 receptors per μm2 on average (Fig. 4A), which is somewhat lower than the 646/μm_2_ reported for A431 cells, a cell line with the highest known EGFR expression level (*52*). This concentration is nevertheless similar to the value observed for the dissociation constant for EGFR-Δ107-fp (Table 1) and indicates that a substantial fraction of EGFR-EYFP is likely to exist in oligomers in these cell lines.

**Figure 4.**
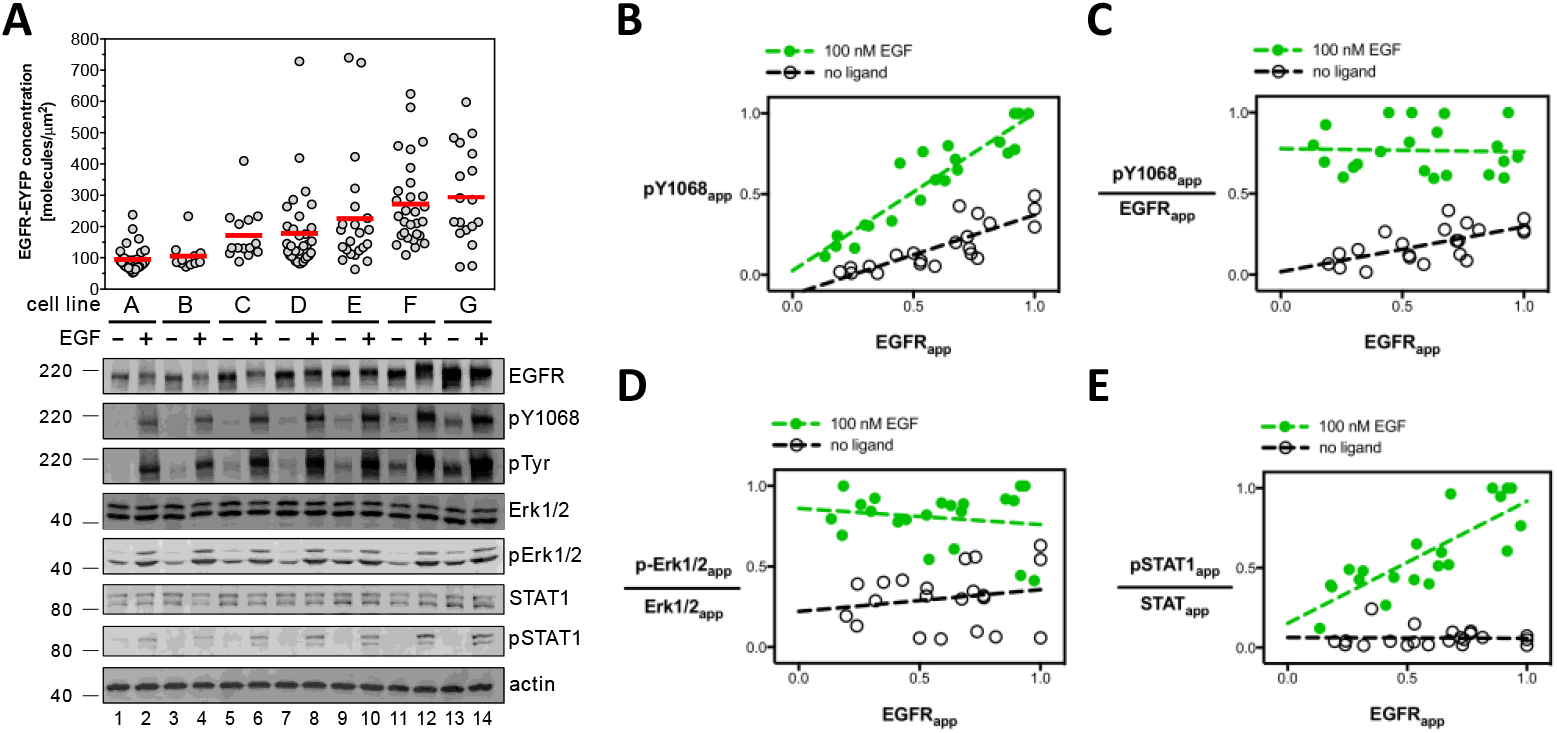
Increased EGFR cell surface concentration leads to increased EGFR phosphorylation but not increased effector phosphorylation in the absence of ligand. (A) Confocal microscopy and western blot analysis of EGF-independent and EGF-dependent EGFR signal transduction. Measurements from seven unique CHO cell lines (cell line A, B, C, ‖, G) are shown, each stably expressing full length EGFR-EYFP. The concentration of EGFR-EYFP as determined by confocal microscopy is plotted on the y-axis. Each grey dot corresponds to a measurement from a single vesicle. Primary antibodies are indicated to the right of the western blots, which are representative of three independent experiments. Molecular weight markers (kDa) indicated to the left of each western blot. Each of the seven stable cell lines (A-G) were treated and analyzed in parallel for a single experiment. Selected western blot groups were quantified in ImageJ and plotted in panels B-E. All values in panels B-E represent integrated band intensities, and the highest value within the experiment for a particular axis was normalized to equal a value of one. Green circles depict bands from wells that were treated with 100 nM EGF, white circles represent bands from wells which were left untreated. Dotted lines show fits to a straight line. (B) Phosphorylated-EGFR (pY-1068) is plotted on the y-axis, total EGFR level on the x-axis. (C) Phosphorylated-EGFR (pY-1068) divided by total EGFR level plotted on the y-axis; total EGFR level on the x-axis. (D) Phospho-Erk1/2 and (E) Phospho-STAT1, divided by the signal for total ERK1/2 and STAT1 proteins, respectively, as a function of total EGFR. Molecular weight markers (kDa) are indicated to the right of each western blot.

Seven cell lines selected for increasing EGFR-EYFP expression levels were treated with EGF and analyzed by western blot for EGFR and phospho-tyrosine (Fig. 4A-C). In the absence of ligand, EGFR phosphorylation was not detectable at concentration levels below 100 molecules/μm2. Between 100 and 300 molecules/μm2, ligand-independent phosphorylation increased in a linear fashion. Stimulation with EGF resulted in a linear increase in phosphorylation across all measured expression levels (Fig. 4C). Notably, the EGFR-EYFP expression range over which EGF-independent phosphorylation begins to become detectable (100-300 molecules/μm2) is comparable to the best fit value for the 2D-dissociation constant for EGFR-Δ107-fp (156 molecules/μm2, Table 1) suggesting that concentration-dependent oligomerization might underlie ligand-independent EGFR phosphorylation.

To determine if EGFR phosphorylation correlated with phosphorylation of downstream effectors in CHO cells, the influence of EGFR membrane concentration on two downstream signaling pathways (STAT and MAPK/Erk) was examined by Western blot (Fig. 4). Increasing EGFR expression had no measurable effect on Erk1/2 phosphorylation or expression in the absence or presence of EGF (Figs. 4A and 4D). Total STAT1 expression levels increased modestly with increasing EGFR expression, but, unlike EGFR, phosphorylation of STAT1 did not increase with increasing EGFR in the absence of ligand. The level of EGF-dependent pSTAT1 was enhanced in EGFR-YFP overexpressing cells, however (Fig. 4E). Thus, although higher EGFR concentrations result in higher basal levels of EGFR phosphorylation, this increase in ligand-independent phosphorylation does not trigger phosphorylation of downstream effectors.

### Asymmetric kinase dimer interface mutations alter but do not abolish ligand-independent EGFR oligomer ensembles

A key element of EGFR activation is formation of an asymmetric dimer interface between the N-lobe of one kinase domain and the C-lobe of another kinase domain. Single amino-acid substitutions at these N-lobe (I706Q) and C-lobe (V948R) interfaces abolish EGF-dependent EGFR activity, but co-expression of these two point mutants restores activity owing to their ability to complement each other’s defect (*37*). To determine if the asymmetric kinase dimer interface is involved in the formation of EGFR oligomers in the absence of ligand, the I706Q and V948R mutations were introduced into the EGFR-Δ107- fp variant and the oligomerization propensity of each variant measured by QI-FRET (Fig. 5). Neither variant had a strong effect on the propensity of EGFR to form ligand-independent oligomers, which contrasts with previous observations that these mutations reduce ligand-dependent oligomers (*41*). In the absence of EGF, the dissociation constants did not significantly differ between the wildtype, I706Q and V948R EGFR-Δ107-fp variants (Fig. 5B-C, Table 1).

**Figure 5.**
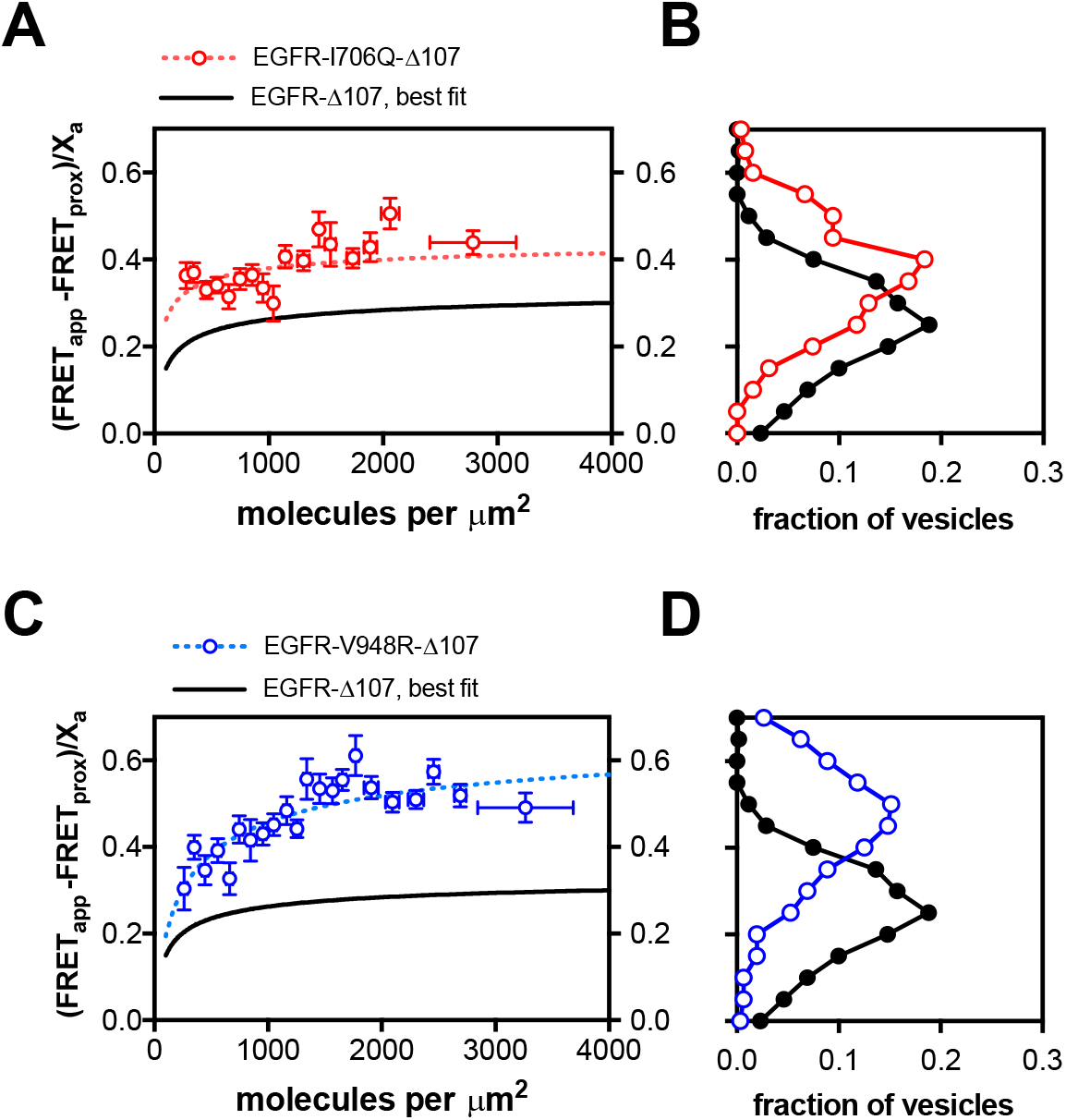
The EGFR missense variants (I706Q and V948R) form ligand-independent oligomers that are structurally distinct from the wildtype oligomer. FRET efficiency for the EGFR-Δ107-fp variants I706Q (A, B) and V948R (C, D). Plots of the FRET efficiency (y-axis, FRET_app_) as a function of receptor concentration (x-axis, units in molecules per square micron) for I706Q (A) and V948R (C). Each protein is indicated above the graph. The black line shows the best fit line for EGFR-Δ107-FP from Figure 2B. The dotted lines represent the best fit line for the I706Q (red) and V948R (blue) variants. As in Figures 2 and 3, the FRET data in 5A and 5C were projected onto the y-axis to yield FRET histograms for EGFR-I706Q-Δ107-fp (B) and EGFR-948R-Δ107-fp (D).

Although these mutations did not alter EGFR oligomerization strength, they did alter the FRET efficiency of the oligomer ensemble. The FRET efficiency in the absence of ligand was much higher for both variants relative to wildtype EGFR-Δ107-fp, with a best-fit parameter for the intrinsic FRET of 0.46 and 0.70 for the I706Q and V948R variants, respectively, vs. 0.35 for EGFR-Δ107-fp (Table 1). The FRET efficiency for both variants was also higher in the presence of ligand relative to wildtype EGFR-Δ107-fp in the presence of ligand (Fig. 5 and Table 1), indicating that disruption of the asymmetric kinase dimer alters the ensemble of intracellular domain interactions in ligand-driven EGFR oligomers.

## Discussion

In this study we use QI-FRET to investigate homo- and hetero-interactions between EGFR and HER2 in vesicles derived from cell membranes. This approach allows measurement of FRET efficiency as a function of receptor concentration owing to variable ErbB levels in vesicles derived from different cells. We observe concentration-dependent FRET efficiency values for several EGFR variants in the absence of ligand, which indicates dynamic equilibria between different EGFR association states and allows derivation of effective 2-dimensional dissociation constants. The homo-oligomerization strength of EGFR observed here, both in the presence and absence of ligand, is similar to previously published results (*23, 24*). Mutations at the EGFR asymmetric kinase dimer interface alter the ensemble of ligand-independent EGFR oligomers but do not significantly alter their stability, implying that ligand-independent EGFR oligomers are distinct from the active state dimer. We also show that although levels of phosphorylated EGFR increase with increasing EGFR concentrations, this phosphorylation does not lead to increased phosphorylation of either Erk1/2 or STAT1, indicating that receptor phosphorylation is not sufficient to trigger pathway activation as recently observed for artificially forced EGFR dimers (*53, 54*).

Understanding the role of ligand-independent EGFR oligomers in regulating ErbB activity remains the topic of much investigation (*18, 19, 41, 55–57*). To estimate the oligomeric fraction of unliganded EGFR and HER2 in physiological membranes using the two-dimensional dissociation constants reported here, the concentration of ERBBs in cell membranes must be known. One recent study reported receptor concentrations between about 100 and 600 molecules/μm_2_ in cell lines derived from human tumors (*52*). ErbB levels are typically reported in units of receptors per cell, however, and conversion of total receptors to a two-dimensional concentration requires estimation of the area of the cell membrane. Assuming a constant receptor density across the entire plasma membrane and a cell diameter of 15 μM, the dissociation constant for EGFR homodimers in the absence of ligand corresponds to ~100,000 EGFR molecules/cell. Our results thus imply that under physiological expression conditions, HER2 is mostly oligomeric and EGFR is 25-50% oligomeric.

The presence of a substantial fraction of phosphorylated oligomeric EGFR in the absence of ligand (*24, 53, 58*) (Fig. 4) raises the question of whether EGFR is sampling the active dimer state, which involves a specific asymmetric dimer interaction between intracellular kinase domains (*37*). Amino-acid substitutions at this interface that impair the ability of EGFR to become phosphorylated in response to ligand and preclude formation of the asymmetric dimer in crystals do not diminish the stability of ligand-independent EGFR oligomers, however, implying the presence of an inactive EGFR oligomer that is distinct from the active dimer. This inactive oligomer appears to depend on the EGFR intracellular domain as deletion of intracellular domain results in loss or decrease of EGFR association as judged by: FRET (*59*), single molecule tracking (*60*), chemical cross-linking (*61*), and loss of negative cooperativity in a saturation binding experiment (*18*). The ErbB extracellular domains must also play a role in mediating or favoring this inactive oligomer: deletion of the extracellular domains leads to constitutive receptor phosphorylation (*35*), and mutations within the interface of the extended ECD dimer destabilize ligand-independent oligomers (*23, 60*). No conserved interaction between the ErbB extracellular domains has been observed in crystals of tethered forms of ErbB ECDs, however, and it is not clear how they inhibit receptor activity in the absence of ligand.

In contrast, a number of interactions observed in crystal structures of inactive ErbB kinase forms have been suggested as reflective of inactive ErbB dimers (*39, 62, 63*). A consensus has yet to emerge on the physiological relevance of these dimers, however, and truncation of ErbB C-terminal tail domains in crystallized forms of inactive ErbB kinases may remove essential components of physiological interactions in the inactive state. Regions of the EGFR C-tail immediately following the kinase domain appear to play a role in stabilizing an inactive state as deletions or mutations in this region lead to enhanced or ligand-independent receptor activity (*29, 64, 65*). An attractive hypothesis is that the kinase distal region interacts with the kinase C-lobe in a fashion that competes with interactions made in the asymmetric kinase dimer interface (*65*). If so, this contact between the kinase and the tail might stabilize the inactive oligomer, which could underlie the observation that the kinase C-lobe mutation (V948R) alters the inactive oligomer ensemble as reflected in an increase in maximal FRET efficiency.

Conversion from inactive to active forms of ERBBs clearly involves coupled rearrangements and interactions between many structural elements. We provide evidence here that in the absence of ligand a substantial fraction of EGFR exists in an inactive oligomeric state that is distinct from the active state and that HER2 is predominantly oligomeric. Future studies are needed to determine the structure and function of this oligomer, whether it is a shared feature among ERBBs, and how it contributes to regulating ErbB activity in normal and disease states.

## Acknowledgements

Lily Raines, Nuala Del Piccolo and Chris King provided technical support with protein purification, cell culture and fluorescence microscopy. Xiaoling Zhang of the Ross Flow Cytometry Core (Johns Hopkins University, School of Medicine) assisted with fluorescence activated cell sorting. POB and DJL were supported by grants from the National Institutes of Health (NIH 5R01GM099321-17) and the Cancer Prevention Research Institute of Texas (CPRIT RR160023). KH was supported by a grant from the National Science Foundation (NSF MCB-1712740).

---

**Figure S1.**
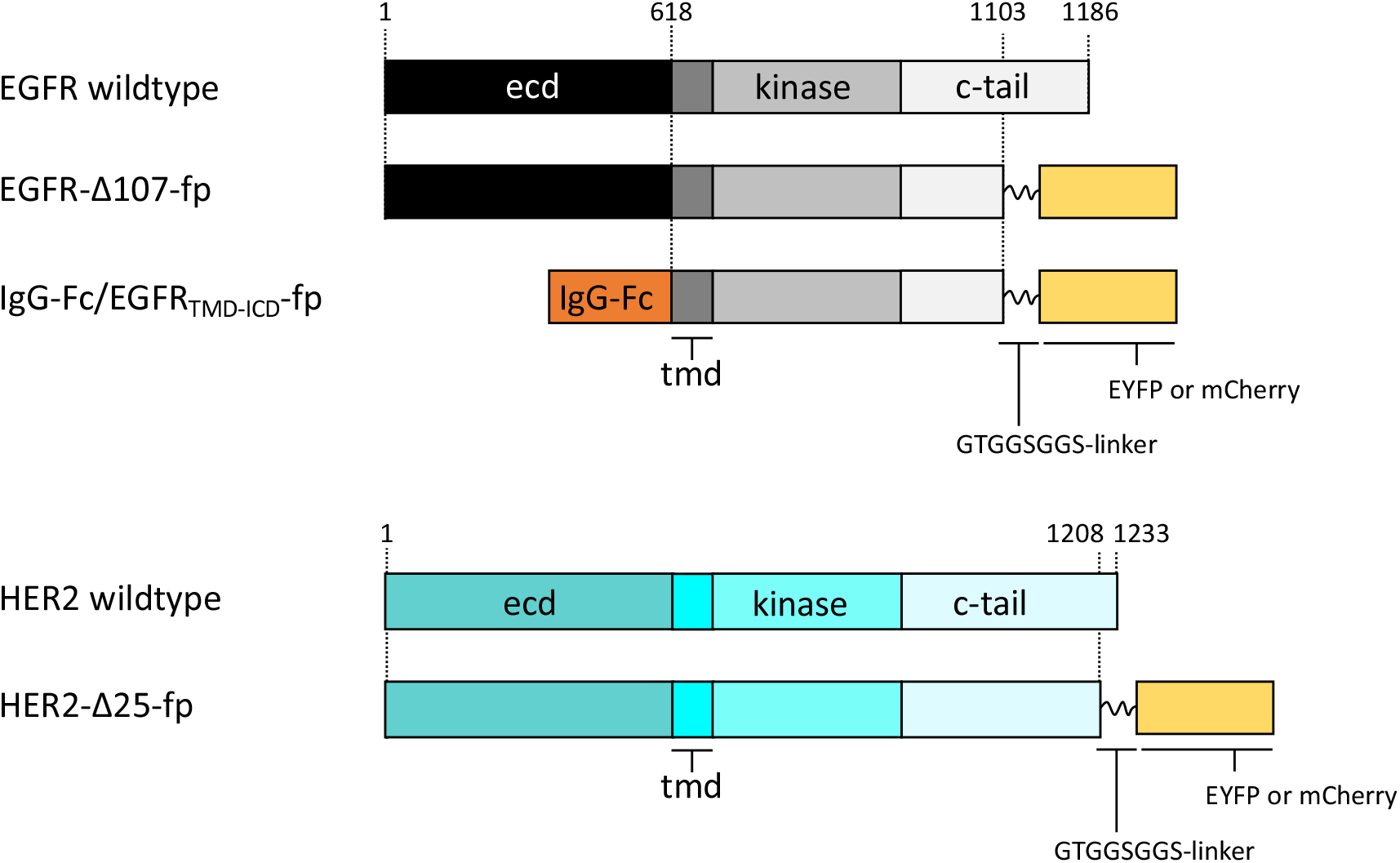
Schematic diagrams of EGFR/ERB1 and HER2/ERBB2 proteins used for FRET studies.

**Supplementary Figure S2.**
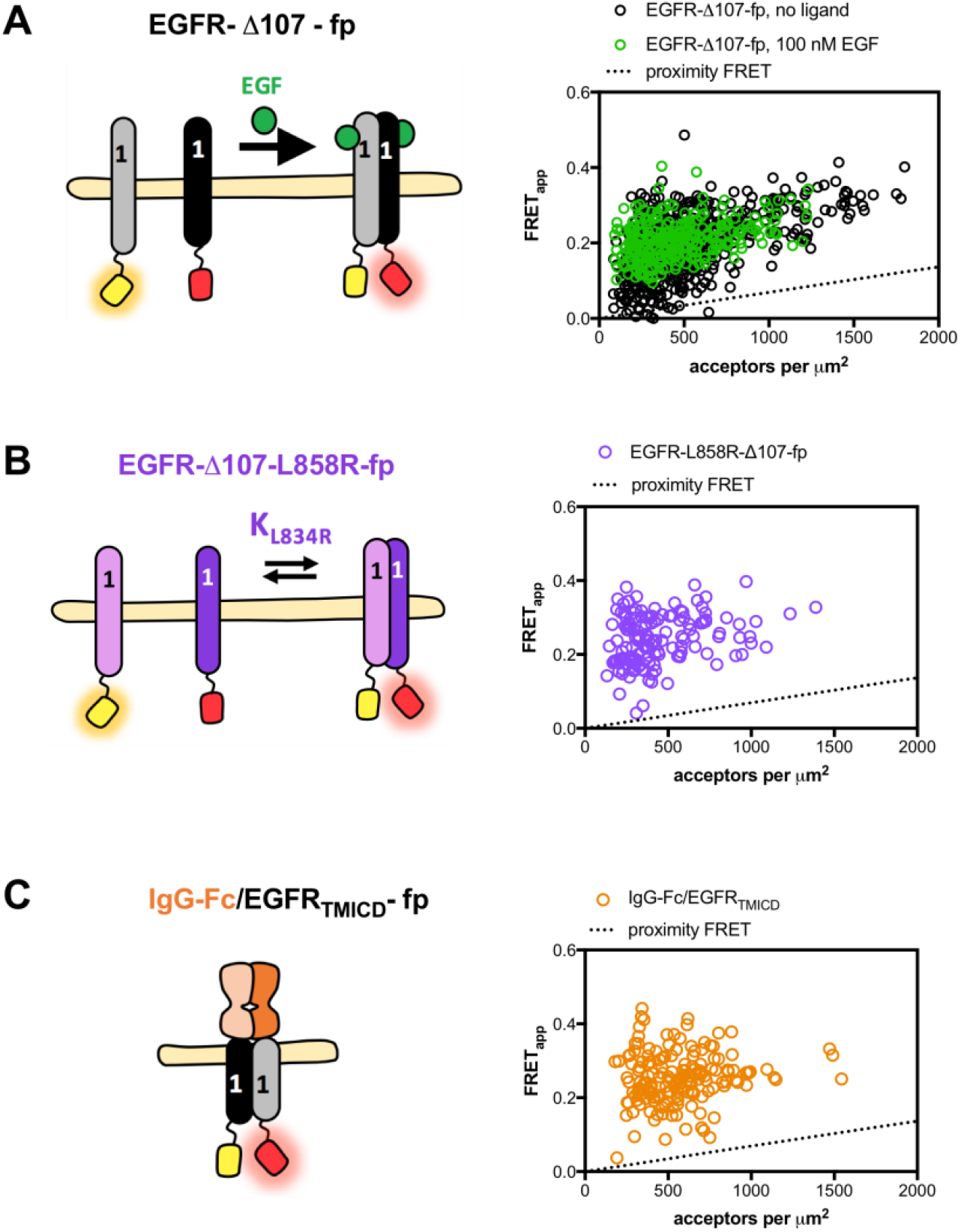
Uncorrected FRET efficiency for EGFR homo-oligomerization. Uncorrected FRET efficiency for EGFR-Δ107-fp wildtype (A), EGFR-Δ107-L858R-fp (B) and IgG-Fc/EGFRTMICD-fp (C). The same data were used to generate these graphs and the graphs in Figure 2. Cartoons are the same as those in Figure 2. Each open circle in the graphs shows the FRET efficiency for a single vesicle. The y-axis shows the apparent FRET efficiency, the x-axis shows the number of acceptor molecules per square micron. The dotted line represents the theoretical FRET efficiency that would occur due to random proximity.

**Supplementary Figure S3.**
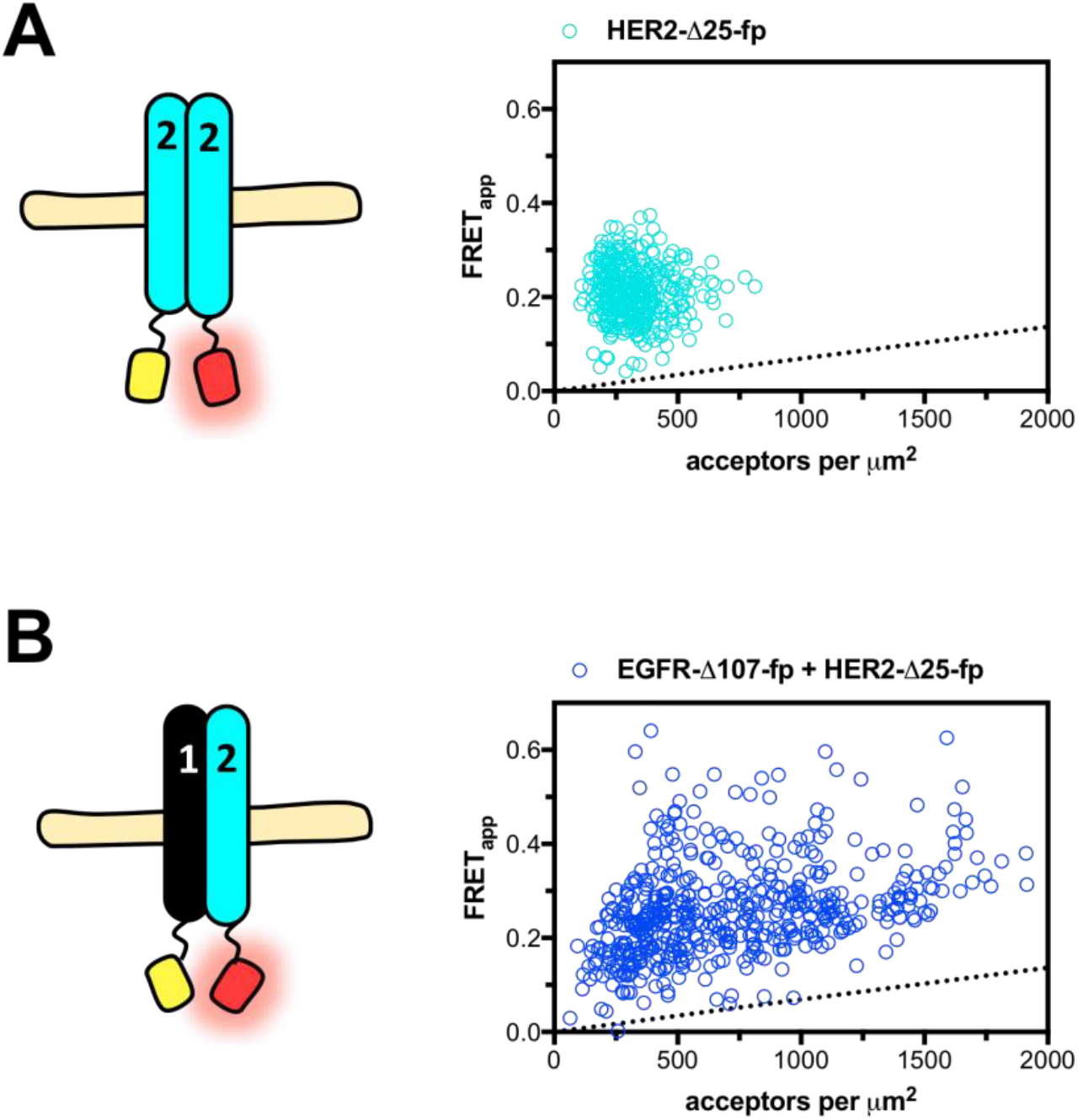
Uncorrected FRET efficiency for HER2 homo- and hetero-oligomerization. Uncorrected FRET efficiency for HER2-Δ25-fp wildtype (A) and (B) HER2-Δ25-fp co-expressed with EGFR-Δ107-fp. The same data were used to generate these graphs and the graphs in Figure 3. Cartoons are the same as those in Figure 3. Each open circle in the graphs shows the FRET efficiency for a single vesicle. The y-axis shows the apparent FRET efficiency, the x-axis shows the number of acceptor molecules per square micron. The dotted line represents the theoretical FRET efficiency that would occur due to random proximity.

**Supplementary Figure S4.**
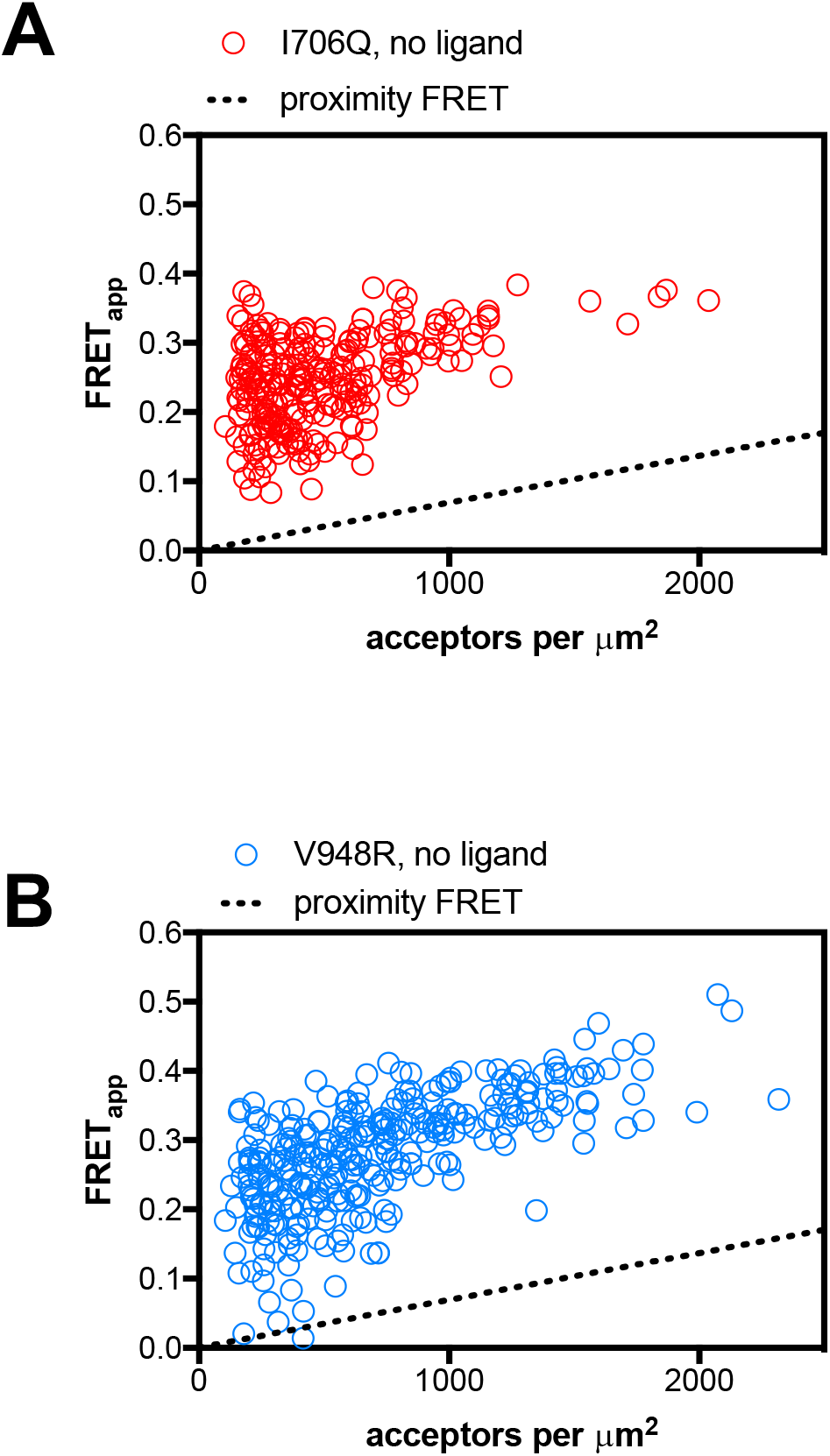
Uncorrected FRET efficiency for the EGFR variants I706Q and V948R. Uncorrected FRET efficiency for the EGFR-Δ107-fp variants I706Q (A) and V948R (B). The same data were used to generate these graphs and the graphs in Figure 5. Each open circle in the graphs shows the FRET efficiency for a single vesicle. The y-axis shows the apparent FRET efficiency, the x-axis shows the number of acceptor molecules per square micron. The dotted line represents the theoretical FRET efficiency that would occur due to random proximity.

**Supplementary Figure S5.**
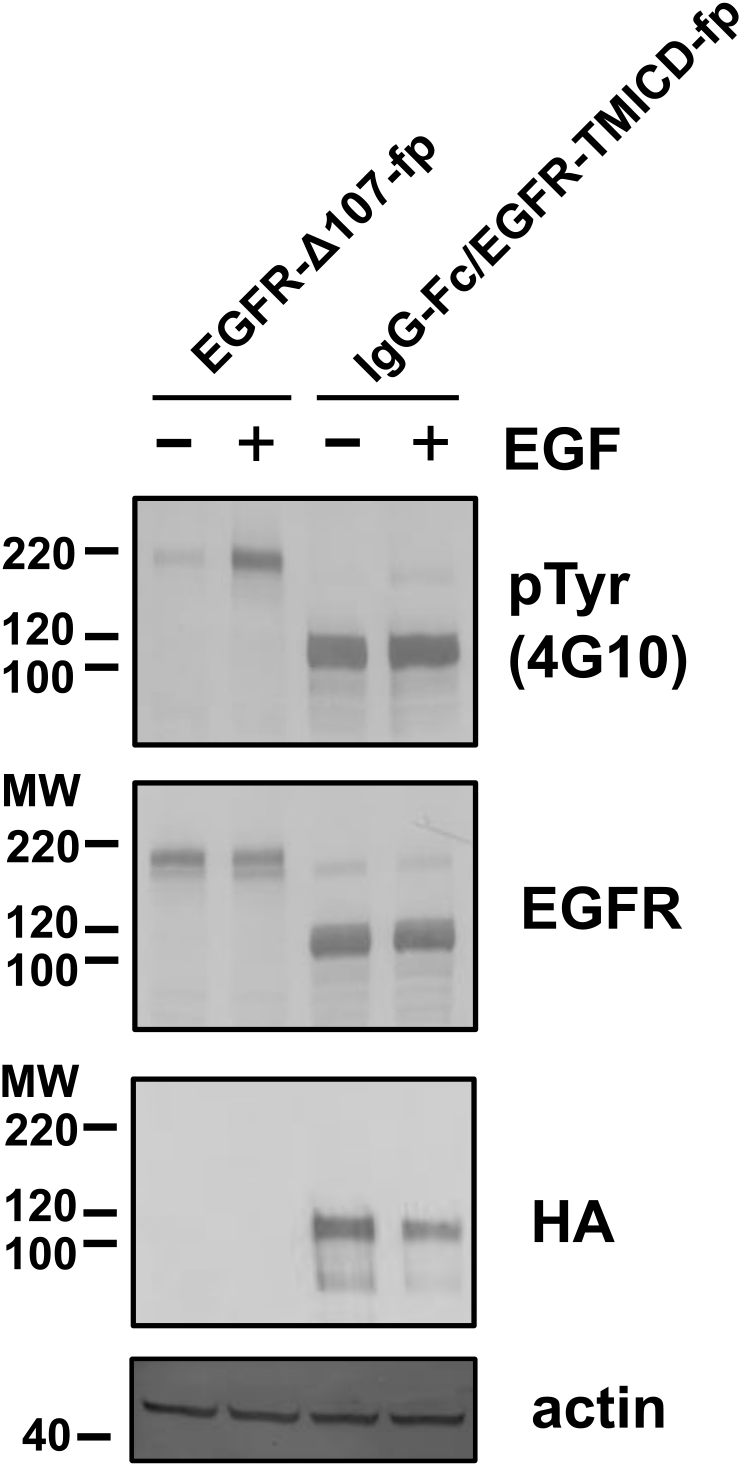
Substitution of the mouse IgG-Fc domain for the EGFR extracellular domain results in a constitutively phosphorylated chimeric receptor. Western blot analysis of CHO cell lysates after transient transfection and treatment with EGF. The indicated plasmid constructs are indicated above the western blots. Antibodies are indicated to the right of each blot, molecular weight markers (kDa) inidicated to the left. The blots are representative of three independent biological replicates.

**Supplementary Table S1.**
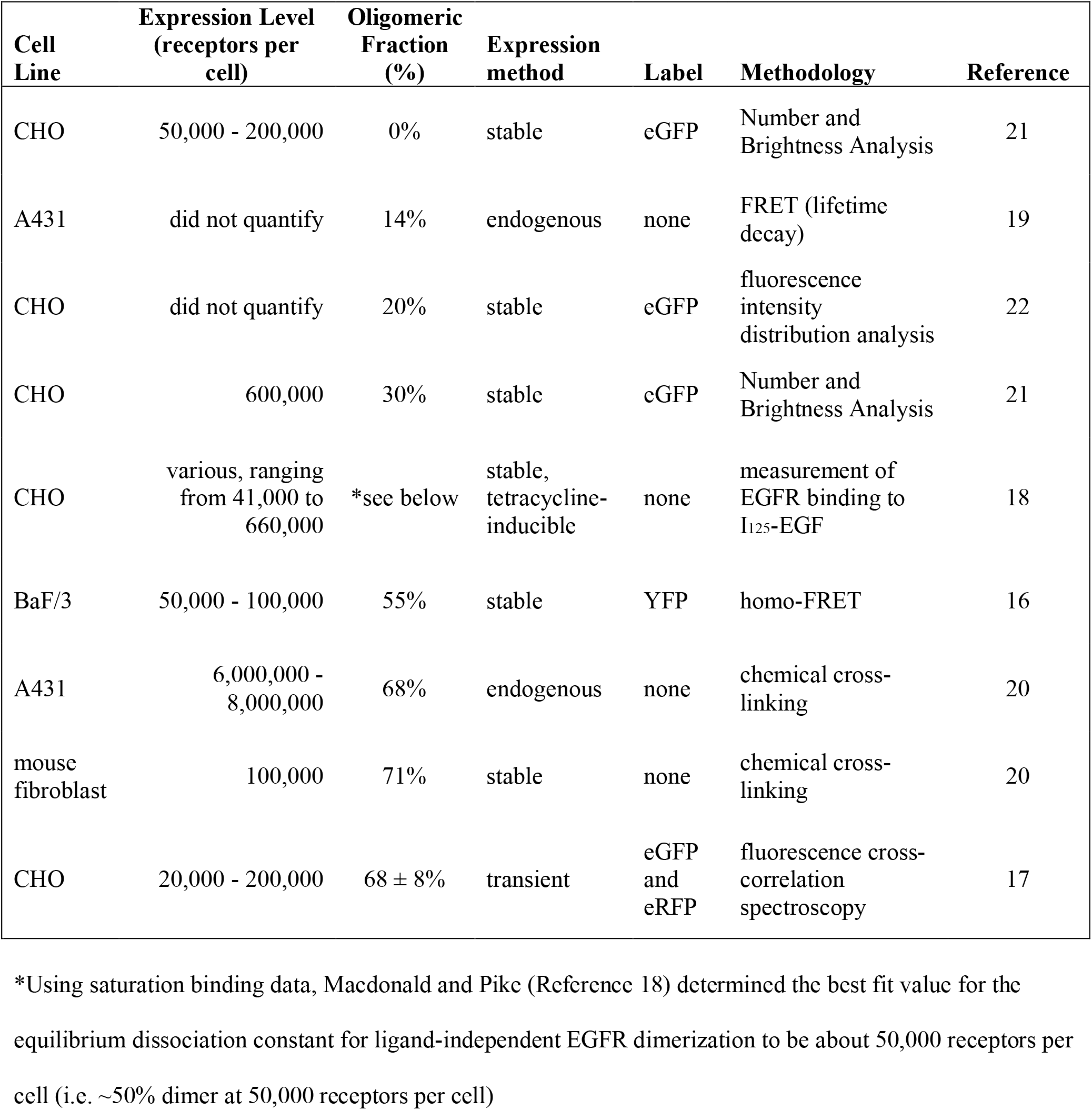
Previously reported fractions of preformed EGFR dimers

| **Cell Line** | **Expression Level**<br>**(receptors per cell)** | **Oligomeric Fraction (%)** | **Expression method** | **Label** | **Methodology** | **Reference** |
| --- | --- | --- | --- | --- | --- | --- |
| CHO | 50,000 - 200,000 | 0% | stable | eGFP | Number and Brightness Analysis | 21 |
| A431 | did not quantify | 14% | endogenous | none | FRET (lifetime decay) | 19 |
| CHO | did not quantify | 20% | stable | eGFP | fluorescence intensity distribution analysis | 22 |
| CHO | 600,000 | 30% | stable | eGFP | Number and Brightness Analysis | 21 |
| CHO | various, ranging from 41,000 to 660,000 | *see below | stable, tetracycline-inducible | none | measurement of EGFR binding to I<sub>125</sub>-EGF | 18 |
| BaF/3 | 50,000 - 100,000 | 55% | stable | YFP | homo-FRET | 16 |
| A431 | 6,000,000 - 8,000,000 | 68% | endogenous | none | chemical cross-linking | 20 |
| mouse fibroblast | 100,000 | 71% | stable | none | chemical cross-linking | 20 |
| CHO | 20,000 - 200,000 | 68 ± 8% | transient | eGFP and eRFP | fluorescence cross-correlation spectroscopy | 17 |

\*Using saturation binding data, Macdonald and Pike (Reference 18) determined the best fit value for the equilibrium dissociation constant for ligand-independent EGFR dimerization to be about 50,000 receptors per cell (i.e. ~50% dimer at 50,000 receptors per cell)

## References

1. M. A. Lemmon, J. Schlessinger, Cell signaling by receptor tyrosine kinases. Cell 141, 1117–1134 (2010).

2. A. G. Sacco, F. P. Worden, Molecularly targeted therapy for the treatment of head and neck cancer: a review of the ErbB family inhibitors. Onco Targets Ther 9, 1927–1943 (2016).

3. N. F. Endres, K. Engel, R. Das, E. Kovacs, J. Kuriyan, Regulation of the catalytic activity of the EGF receptor. Curr Opin Struct Biol 21, 777–784 (2011).

4. Y. Yarden, J. Schlessinger, Epidermal growth factor induces rapid, reversible aggregation of the purified epidermal growth factor receptor. Biochemistry 26, 1443–1451 (1987).

5. Y. Yarden, J. Schlessinger, Self-phosphorylation of epidermal growth factor receptor: evidence for a model of intermolecular allosteric activation. Biochemistry 26, 1434–1442 (1987).

6. G. J. Todaro, J. E. De Larco, S. Cohen, Transformation by murine and feline sarcoma viruses specifically blocks binding of epidermal growth factor to cells. Nature 264, 26–31 (1976).

7. R. N. Fabricant, J. E. De Larco, G. J. Todaro, Nerve growth factor receptors on human melanoma cells in culture. Proc Natl Acad Sci US A 74, 565–569 (1977).

8. R. W. Holley, R. Armour, J. H. Baldwin, K. D. Brown, Y. C. Yeh, Density-dependent regulation of growth of BSC-1 cells in cell culture: control of growth by serum factors. Proc Natl Acad Sci USA 74, 5046–5050 (1977).

9. G. Carpenter, S. Cohen, Biological and molecular studies of the mitogenic effects of human epidermal growth factor. Symp Soc Dev Biol, 13–31 (1978).

10. A. R. Rees, E. D. Adamson, C. F. Graham, Epidermal growth factor receptors increase during the differentiation of embryonal carcinoma cells. Nature 281, 309–311 (1979).

11. M. R. Green, D. A. Basketter, J. R. Couchman, D. A. Rees, Distribution and number of epidermal growth factor receptors in skin is related to epithelial cell growth. Dev Biol 100, 506–512 (1983).

12. J. Filmus, M. N. Pollak, R. Cailleau, R. N. Buick, MDA-468, a human breast cancer cell line with a high number of epidermal growth factor (EGF) receptors, has an amplified EGF receptor gene and is growth inhibited by EGF. Biochem Biophys Res Commun 128, 898–905 (1985).

13. G. P. Cowley, J. A. Smith, B. A. Gusterson, Increased EGF receptors on human squamous carcinoma cell lines. Br J Cancer 53, 223–229 (1986).

14. N. E. Davidson, E. P. Gelmann, M. E. Lippman, R. B. Dickson, Epidermal growth factor receptor gene expression in estrogen receptor-positive and negative human breast cancer cell lines. Mol Endocrinol 1, 216–223 (1987).

15. P. P. Di Fiore et al., Overexpression of the human EGF receptor confers an EGF-dependent transformed phenotype to NIH 3T3 cells. Cell 51, 1063–1070 (1987).

16. N. Kozer et al., Evidence for extended YFP-EGFR dimers in the absence of ligand on the surface of living cells. Phys Biol 8, 066002 (2011).

17. P. Liu et al., Investigation of the dimerization of proteins from the epidermal growth factor receptor family by single wavelength fluorescence cross-correlation spectroscopy. Biophys J 93, 684–698 (2007).

18. J. L. Macdonald, L. J. Pike, Heterogeneity in EGF-binding affinities arises from negative cooperativity in an aggregating system. Proc Natl Acad Sci U S A 105, 112–117 (2008).

19. M. Martin-Fernandez, D. T. Clarke, M. J. Tobin, S. V. Jones, G. R. Jones, Preformed oligomeric epidermal growth factor receptors undergo an ectodomain structure change during signaling. Biophys J 82, 2415–2427 (2002).

20. T. Moriki, H. Maruyama, I. N. Maruyama, Activation of preformed EGF receptor dimers by ligand-induced rotation of the transmembrane domain. J Mol Biol 311, 1011–1026 (2001).

21. P. Nagy, J. Claus, T. M. Jovin, D. J. Arndt-Jovin, Distribution of resting and ligand-bound ErbB1 and ErbB2 receptor tyrosine kinases in living cells using number and brightness analysis. Proc Natl Acad Sci U S A 107, 16524–16529 (2010).

22. S. Saffarian, Y. Li, E. L. Elson, L. J. Pike, Oligomerization of the EGF receptor investigated by live cell fluorescence intensity distribution analysis. Biophys J 93, 1021–1031 (2007).

23. M. R. Stoneman et al., A general method to quantify ligand-driven oligomerization from fluorescence-based images. Nat Methods 16, 493–496 (2019).

24. D. H. Kim et al., Direct visualization of single-molecule membrane protein interactions in living cells. PLoS Biol 16, e2006660 (2018).

25. T. J. Lynch et al., Activating mutations in the epidermal growth factor receptor underlying responsiveness of non-small-cell lung cancer to gefitinib. N Engl J Med 350, 2129–2139 (2004).

26. E. Padfield, H. P. Ellis, K. M. Kurian, Current Therapeutic Advances Targeting EGFR and EGFRvIII in Glioblastoma. Front Oncol 5, 5 (2015).

27. K. B. Pahuja et al., Actionable Activating Oncogenic ERBB2/HER2 Transmembrane and Juxtamembrane Domain Mutations. Cancer Cell 34, 792–806 e795 (2018).

28. E. Penuel, R. W. Akita, M. X. Sliwkowski, Identification of a region within the ErbB2/HER2 intracellular domain that is necessary for ligand-independent association. J Biol Chem 277, 28468–28473 (2002).

29. G. Pines, P. H. Huang, Y. Zwang, F. M. White, Y. Yarden, EGFRvIV: a previously uncharacterized oncogenic mutant reveals a kinase autoinhibitory mechanism. Oncogene 29, 5850–5860 (2010).

30. P. Stephens et al., Lung cancer: intragenic ERBB2 kinase mutations in tumours. Nature 431, 525–526 (2004).

31. A. W. Burgess et al., An open-and-shut case? Recent insights into the activation of EGF/ErbB receptors. Mol Cell 12, 541–552 (2003).

32. D. Diwanji, T. Thaker, N. Jura, More than the sum of the parts: Toward full-length receptor tyrosine kinase structures. IUBMB Life 71, 706–720 (2019).

33. S. Bouyain, P. A. Longo, S. Li, K. M. Ferguson, D. J. Leahy, The extracellular region of ErbB4 adopts a tethered conformation in the absence of ligand. Proc Natl Acad Sci U S A 102, 15024–15029 (2005).

34. H. S. Cho, D. J. Leahy, Structure of the extracellular region of HER3 reveals an interdomain tether. Science 297, 1330–1333 (2002).

35. N. F. Endres et al., Conformational coupling across the plasma membrane in activation of the EGF receptor. Cell 152, 543–556 (2013).

36. K. M. Ferguson et al., EGF activates its receptor by removing interactions that autoinhibit ectodomain dimerization. Mol Cell 11, 507–517 (2003).

37. X. Zhang, J. Gureasko, K. Shen, P. A. Cole, J. Kuriyan, An allosteric mechanism for activation of the kinase domain of epidermal growth factor receptor. Cell 125, 1137–1149 (2006).

38. A. Arkhipov et al., Architecture and membrane interactions of the EGF receptor. Cell 152, 557–569 (2013).

39. M. Landau, S. J. Fleishman, N. Ben-Tal, A putative mechanism for downregulation of the catalytic activity of the EGF receptor via direct contact between its kinase and C-terminal domains. Structure 12, 2265–2275 (2004).

40. T. S. Wehrman et al., A system for quantifying dynamic protein interactions defines a role for Herceptin in modulating ErbB2 interactions. Proc Natl Acad Sci U S A 103, 19063–19068 (2006).

41. Y. Huang et al., Molecular basis for multimerization in the activation of the epidermal growth factor receptor. Elife 5, (2016).

42. J. M. Kavran et al., How IGF-1 activates its receptor. Elife 3, (2014).

43. S. Sarabipour, R. B. Chan, B. Zhou, G. Di Paolo, K. Hristova, Analytical characterization of plasma membrane-derived vesicles produced via osmotic and chemical vesiculation. Biochim Biophys Acta 1848, 1591–1598 (2015).

44. L. Chen, L. Novicky, M. Merzlyakov, T. Hristov, K. Hristova, Measuring the energetics of membrane protein dimerization in mammalian membranes. J Am Chem Soc 132, 3628–3635 (2010).

45. L. Chen, J. Placone, L. Novicky, K. Hristova, The extracellular domain of fibroblast growth factor receptor 3 inhibits ligand-independent dimerization. Sci Signal 3, ra86 (2010).

46. C. Qiu et al., In vitro enzymatic characterization of near full length EGFR in activated and inhibited states. Biochemistry 48, 6624–6632 (2009).

47. T. H. Evers, E. M. van Dongen, A. C. Faesen, E. W. Meijer, M. Merkx, Quantitative understanding of the energy transfer between fluorescent proteins connected via flexible peptide linkers. Biochemistry 45, 13183–13192 (2006).

48. E. Li, J. Placone, M. Merzlyakov, K. Hristova, Quantitative measurements of protein interactions in a crowded cellular environment. Anal Chem 80, 5976–5985 (2008).

49. P. K. Wolber, B. S. Hudson, An analytic solution to the Forster energy transfer problem in two dimensions. Biophys J 28, 197–210 (1979).

50. C. King, V. Raicu, K. Hristova, Understanding the FRET Signatures of Interacting Membrane Proteins. J Biol Chem 292, 5291–5310 (2017).

51. C. C. Valley et al., Enhanced dimerization drives ligand-independent activity of mutant epidermal growth factor receptor in lung cancer. Mol Biol Cell 26, 4087–4099 (2015).

52. F. Zhang et al., Quantification of epidermal growth factor receptor expression level and binding kinetics on cell surfaces by surface plasmon resonance imaging. Anal Chem 87, 9960–9965 (2015).

53. S. I. Liang et al., Phosphorylated EGFR Dimers Are Not Sufficient to Activate Ras. Cell Rep 22, 2593–2600 (2018).

54. T. Yoshida et al., Matuzumab and cetuximab activate the epidermal growth factor receptor but fail to trigger downstream signaling by Akt or Erk. Int J Cancer 122, 1530–1538 (2008).

55. C. W. Ward, M. C. Lawrence, V. A. Streltsov, T. E. Adams, N. M. McKern, The insulin and EGF receptor structures: new insights into ligand-induced receptor activation. Trends Biochem Sci 32, 129–137 (2007).

56. J. L. Macdonald-Obermann, S. Adak, R. Landgraf, D. Piwnica-Worms, L. J. Pike, Dynamic analysis of the epidermal growth factor (EGF) receptor-ErbB2-ErbB3 protein network by luciferase fragment complementation imaging. J Biol Chem 288, 30773–30784 (2013).

57. E. R. Purba, E. I. Saita, I. N. Maruyama, Activation of the EGF Receptor by Ligand Binding and Oncogenic Mutations: The “Rotation Model”. Cells 6, (2017).

58. C. King, M. Stoneman, V. Raicu, K. Hristova, Fully quantified spectral imaging reveals in vivo membrane protein interactions. Integr Biol (Camb) 8, 216–229 (2016).

59. D. M. Freed et al., EGFR Ligands Differentially Stabilize Receptor Dimers to Specify Signaling Kinetics. Cell 171, 683–695 e618 (2017).

60. I. Chung et al., Spatial control of EGF receptor activation by reversible dimerization on living cells. Nature 464, 783–787 (2010).

61. X. Yu, K. D. Sharma, T. Takahashi, R. Iwamoto, E. Mekada, Ligand-independent dimer formation of epidermal growth factor receptor (EGFR) is a step separable from ligand-induced EGFR signaling. Mol Biol Cell 13, 2547–2557 (2002).

62. N. Jura et al., Mechanism for activation of the EGF receptor catalytic domain by the juxtamembrane segment. Cell 137, 1293–1307 (2009).

63. N. Jura, Y. Shan, X. Cao, D. E. Shaw, J. Kuriyan, Structural analysis of the catalytically inactive kinase domain of the human EGF receptor 3. Proc Natl Acad Sci U S A 106, 21608–21613 (2009).

64. E. Kovacs et al., Analysis of the Role of the C-Terminal Tail in the Regulation of the Epidermal Growth Factor Receptor. Mol Cell Biol 35, 3083–3102 (2015).

65. R. Y. Yang, K. S. Yang, L. J. Pike, G. R. Marshall, Targeting the dimerization of epidermal growth factor receptors with small-molecule inhibitors. Chem Biol Drug Des 76, 1–9 (2010).

